# Gene expression pattern of vacuolar-iron transporter-like (VTL) genes in hexaploid wheat during metal stress

**DOI:** 10.1101/863084

**Authors:** Shivani Sharma, Gazaldeep Kaur, Anil Kumar, Varsha Meena, Hasthi Ram, Jaspreet Kaur, Ajay Kumar Pandey

**Author notes:** Corresponding author, Dr. Ajay K Pandey, Scientist-E, National Agri-Food Biotechnology Institute (Department of Biotechnology), Sector 81, Knowledge City, Mohali-140306, Punjab, India.

## Abstract

Iron is one of the important micronutrients that is not just essential for the human body, but also required for crop productivity and yield-related traits. To address the Fe homeostasis in crop plants, multiple transporters belonging to the category of Major facilitator superfamily are being explored. In this direction, Vacuolar iron transporters (VIT) are being reported and have been characterized functionally as an important candidate to address biofortification in cereal crops. In the present study, the identification and characterization of new members of <u>V</u>acuolar iron <u>t</u>ransporters-<u>l</u>ike proteins (VTL) was performed. Phylogenetic analyses demonstrated distinct clustering of all the *VTL* genes from the previously known *VIT* genes. Our analysis identifies multiple *VTL* genes from hexaploid wheat with the highest number of this gene family localized on chromosome 2. Quantitative expression analysis suggests that most of the *VTL* genes are induced only during the Fe surplus condition, thereby reinforcing their role metal homeostasis. Interestingly, most of the wheat *VTL* genes were significantly up-regulated in a tissue-specific manner under Zn, Mn and Cu deficiency conditions. Although, no significant changes in expression of wheat *VTL* genes were observed in roots under heavy metals, but *TaVTL2*, *TaVTL3* and *TaVTL5* were upregulated in the presence of cobalt stress. Overall, this work deals with the characterization of wheat *VTL* genes that could provide an important genetic resource for addressing metal homeostasis in bread wheat.

## 1. Introduction

Successful micronutrient biofortification of crops through biotechnology requires detailed knowledge of complex homeostatic mechanisms that tightly regulate the micronutrient concentrations in plants. Iron (Fe) is one of the important micronutrients that is involved in multiple important cellular and physiological processes in plants [1–3]. Some of the important functions include its importance in photosynthesis, nitrogen fixation and respiration [4,5]. Although Fe may be present in the soil, yet due to alkaline rhizospheric conditions or unfavorable circumstances it is not been efficiently taken up by plants [6–9]. Moreover, the Fe is mobilized inside the plant tissue with an important goal to load in the filial tissue of grains that basically involves a multistep process encompassing many bottlenecks [10–13]. Researchers worldwide is generating information for the means to enrich Fe rich grains and their storage with enhanced bioavailability. To improve Fe content in cereal grains, multiple transporters and chelators have been targeted including molecular approaches [14–16]. In addition to these, a number of genes are still unaddressed that are potential candidates for micronutrient biofortification, including transporters belong to the inventory of the Major facilitator superfamily (MFS) gene family [17]. Also, a very few reports are available that deals with the identification and molecular characterization of wheat genes or gene families involved in Fe and Zn homeostasis. Recent reports are emerging for the identification of few functional gene families, for example belonging to, yellow stripe like transporters [18], nicotianamine synthase (NAS), deoxymugineic Acid Synthase (DMAS) [19], yet there are still many genes families that remained to be characterized in hexaploid wheat. Similarly, genes encoding for Zinc–Induced Facilitator-Like family (ZIFL) of transporters have been described for their role during Fe homeostasis beside been regulated by a few heavy metals [20].

Fe storage in seeds gets compartmentalized in two major subcellular stores that include chloroplasts and vacuoles. For example, 95% of the iron is stored in vacuoles in the *Arabidopsis* seeds [21]. Vacuoles are an important site for Fe mobilization wherein, they are bound to various chelators like phytic acid, nicotianamine and other organic acids etc. Therefore, uptake of Fe into vacuoles could be an alternate strategy to enhance total micronutrient content with a minimized tradeoff for its toxicity in the tissue. To design such strategy, the role of vacuolar transporters needs to be addressed and exploited [15,22].

Previously, one such vacuolar iron transporters (VIT) were shown to be playing an important role to maintain Fe in the optimal physiological range and prevent cellular toxicity. *VIT* genes from multiple plant species have been characterized and assessed for their ability to enhance Fe content in cereal crops [15]. These *VIT* genes show high homology with a small family of nodulin like protein containing a CCC-1 (Ca^2+^-Sensitive Cross Complementer) like domain with yeast Cccp1 [23]. CCC-1 like the domain was initially discovered in yeast encoded for the vacuolar iron transporter in yeast. Furthermore, mutant *ccc1* cells show increased sensitivity to external iron [21,24] *AtVIT1* is one of the early characterized genes showing the presence of CCC-1 like domain and transport of iron to vacuoles [21]. Utilizing the bioinformatics resources, subsequent studies led to the identification of many <u>v</u>acuolar iron <u>t</u>ransporters-<u>l</u>ike (VTL) proteins from different plant species. Model species, *Arabidopsis* genome encodes five VTL proteins and overexpression of the few genes have shown increased Fe content in seeds. AtVIT1 protein can transport iron into the vacuoles to counter the toxicity and support the seedling development under enhanced iron conditions [23,25]. This suggested that these genes likely to have a function in regulation in Fe homeostasis.

Therefore, the characterization of vacuolar transporters in an important crop such as wheat becomes a prerequisite to address the global issue of biofortification. Wheat is an important crop that is consumed in many developing countries, including India and is therefore being targeted for trait improvement for nutritional quality. In the current work genome-wide identification of wheat *VTL* genes was performed. Further, expression studies during different regimes of Fe, Zn and heavy metals was done that provide an insight into the regulation of wheat *VTL* genes in a tissue-specific manner.

## 2. Results

### 2.1 Identification, phylogenetic analysis and genomic distribution of wheat VTL genes

Thirty-one wheat VIT family sequences were identified based on Ensembl Pfam search and bidirectional blast analysis (Table S1). Subsequently, to study the phylogenetic relationship among VIT family protein sequences from wheat, *Brachypodium*, maize, rice, *Arabidopsis* and *S. cerevisiae*, an unrooted neighbour-joining tree was constructed. This analysis separated the sequences into two distinct clades for VTL and VIT proteins that corresponds to the previously stated distribution of the VIT family. This also led to the classification of the wheat VIT family into 8 VIT and 23 VTL sequences (Figure 1, Table S2). Due to the occurrence of homoeologs, the 23 VTL sequences representing only 4 *VTL* genes, named as *TaVTL1*, *TaVTL2*, *TaVTL4* and *TaVTL5* based on their corresponding rice orthologs followed by the chromosome on which they are present. None of the orthologs in wheat showed high confidence similarity with rice vacuolar iron transporter homolog 3. *TaVTL1* and *4* were found to have three homoeologs, while *TaVTL2* has 4. In contrast, the phylogenetic analysis grouped 13 highly similar sequences together with rice vacuolar iron transporter homolog 5, these were named as *TaVTL5* (Figure S1).

**Figure 1:**
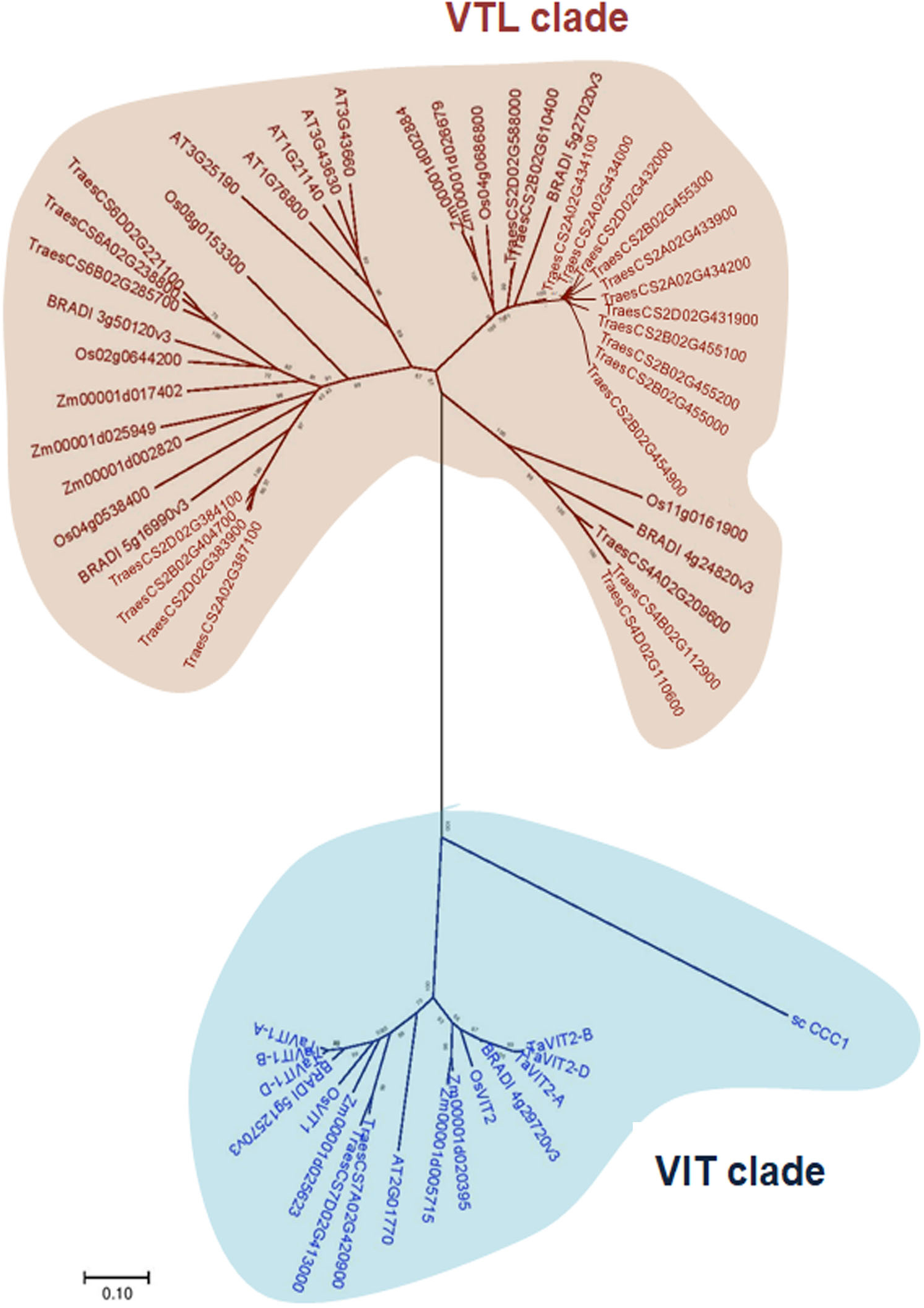
Phylogenetic analysis showing separation of VIT family genes in *Arabidopsis*, *Brachypodium*, *Oryza sativa*, *Zea mays* and *Triticum aestivum* into two distinct clades; VTL clade and VIT clade. The Neighbour-joining phylogenetic tree was generated using MEGA. The numbers represent bootstrap values from 1000 replicates.

*TaVIT1* and *TaVIT2* have already been reported earlier [15]. Interestingly, another new wheat *VIT* with two homoeologs at chromosome 7 (sub-genomes A and D) was identified and referred as *TaVIT3*. *VIT* genes are located on chromosome groups 2, 5 and 7 while *VTL* genes on chromosome groups 2, 4, and 6 with maximum contribution from chromosome 2. Nine *VTL* genes are present on B sub-genome, while seven each on A and D sub-genomes. Maximum number of VTL sequences are located on chromosome 2B (Figure 2A)._

**Figure 2:**
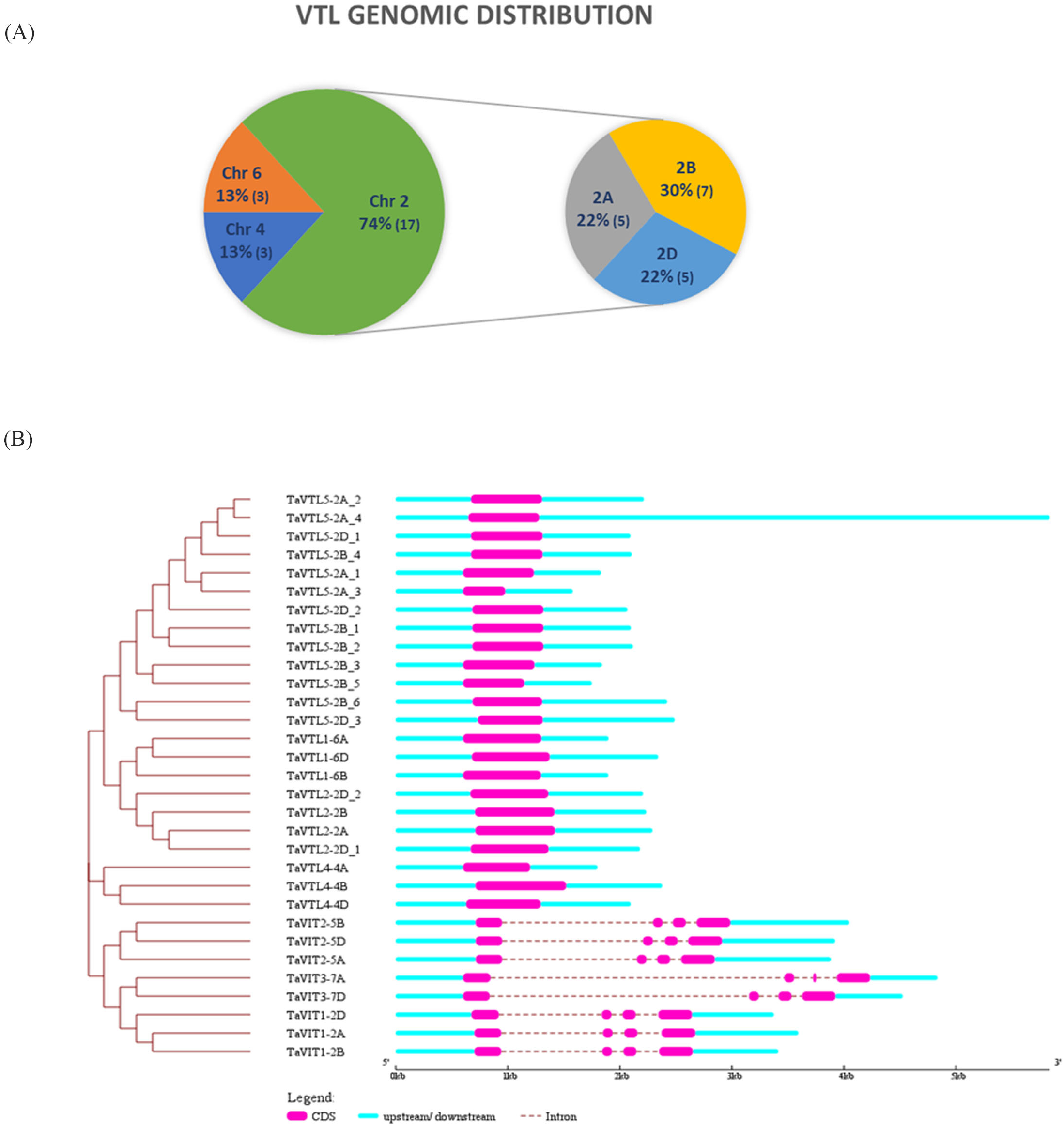
Genomic distribution and exon-intron arrangements of VTL genes. (A) Wheat VTL genes were present on chromosome groups 2, 4 and 6 with maximum VTL genes on chromosome group 2, which was selected to show the VTL gene distribution on 2A, 2B and 2D chromosomes. (B) Genomic structure for wheat *VTL* and *VIT* genes. The intron-exon arrangement was identified using Gene Structure Display Server (GSDS). Exons and introns are represented using pink boxes and cyan lines, respectively. The scale determines the size of the genomic regions.

### 2.2 Gene, protein structure and subcellular localization

*VIT* genes in wheat have three and four intronic and exonic regions respectively, while *VTL* genes have a single exon each with the absence of any introns (Figure 2B), clearly dividing the VIT family into two sub-families based on gene structure also. CDS length was found to be varying from 657 to 747 nucleotides for wheat *VIT* genes. The CDS length for *VTL* genes was ranging from 549 to 810 nucleotides except for *TaVTL5-2A_3* that was 378 nucleotides long. The short length of one VTL gene is due to the missing sequence information at the stop site. The length of TaVIT peptides ranged from 218 to 256 while TaVTL protein length varied from 125 to 269 amino acids. The division of VIT and VTL proteins was also evident from the sub-cellular localization (Table S2); while TaVIT proteins were predicted to be predominantly localized on the plasma membrane and chloroplast thylakoid membrane, maximum TaVTL proteins were predicted to be present on the vacuolar membrane (87%). TaVTL4-4A was predicted to be localized on plasma membrane. VIT proteins had 3-4 predicted TM domains. TaVTL1, 2 and 4 have 5 TM domains majorly, except for TaVTL4-4B which was predicted to have 6 TM domains. Only TaVTL5-2D_3 had 5 TM domains; other paralogs/homoeologs of TaVTL5 had lesser number of TM domains probably due to gene duplication events or missing information. To summarize, TaVTLs have 5 TM domains predominantly, which are depicted in Table S2. Eucalyptus VIT1 (EgVIT1) crystal structure was deciphered recently [26] that was used to confirm the VIT family protein topology prediction using Phobius [27]. EgVIT1 was predicted to have only three TM domains while the crystal structure stated the presence of five TM domains. Therefore, VIT, as well as VTL protein sequences from wheat, were aligned to EgVIT1 to see the possible TM domains in addition to those predicted by Phobius (Figure S2).

### 2.3 Conserved domain and motif analysis

All the VIT and *VTL* genes were found to have the typical CCC1-like superfamily domains of yeast, which were demonstrated earlier for the iron and manganese transport from the cytosol to vacuole. Motif analysis using MEME webserver suggested that motifs 6, 9, and 10 are VIT specific with exceptions for motif 10 been absent in TaVIT3 and motif 6 absent in VIT3-7A. Similarly, motifs 5, 7, 8, 11 to 14 are VTL specific with the exceptions for motif 5 been absent in TaVTL5-2B_3 and TaVTL5-2B_5, where motif 7 was specific for TaVTL5 sequences except in TaVTL5-2A_3, TaVTL5-2B_6, TaVTL5-2D_3. Motif 8 was specific for TaVTL1, 2 and 4. Motif 11 for TaVTL1 and 2. Motif 12 and 13 were unique for TaVTL2, whereas Motif 14 was present only in TaVTL1 (Figure 3, Table S3).

**Figure 3:**
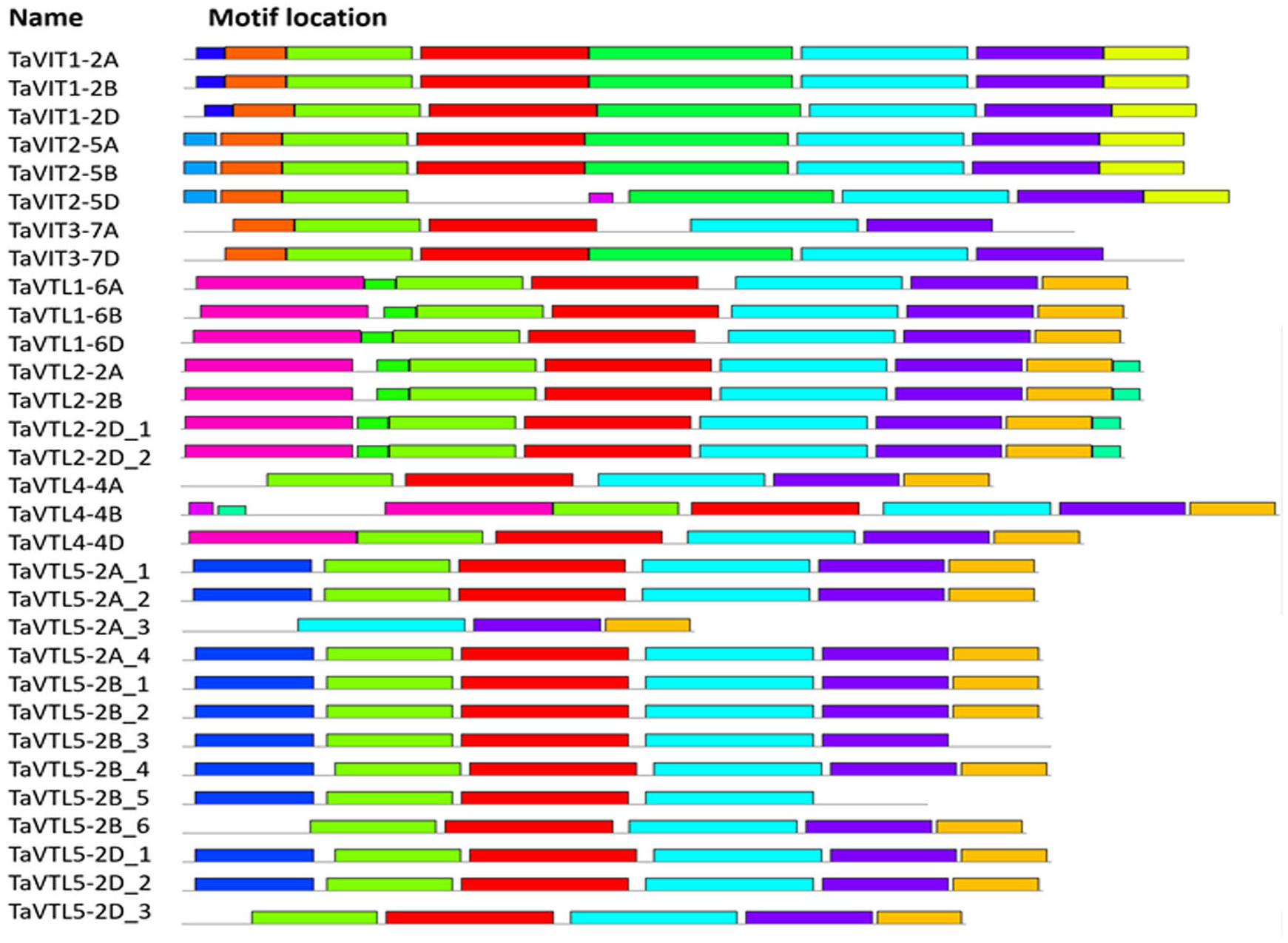
Conserved motifs identified for TaVIT and TaVTL proteins using MEME suite 5.1.0. The colored rectangles on each sequence represent specific conserved motifs numbered 1 through 14, as depicted by the color codes in the box.

### 2.4 Expression of wheat VTL genes under Fe deficiency and surplus condition

To check the regulation of *VTL* genes at the transcriptional level, the promoter for the wheat *VTL* genes were scanned for the cis-elements responsive for Fe and heavy metals. The analysis revealed the presence of multiple such sequences, including <u>i</u>ron-<u>d</u>eficiency-responsive <u>e</u>lement <u>1</u> (IDE1), metal response element (MRE), heavy metal responsive element (HMRE) and iron-related bHLH transcription factor 2 (IRO2) binding site (Table S4). In the most abundant category, <u>i</u>ron-<u>d</u>eficiency-responsive <u>e</u>lement <u>1</u> (IDE1) was predominant. Interestingly, the IRO2 binding site was present only in the regulatory region of *TaVTL2B/D*. These observations suggest that VTL expression could be regulated by the presence-absence of specific metals including micronutrients such as Fe and Zn.

Previously, *VTL* genes were reported to have differential expression patterns under the changing regimes of Fe and Zn [25]. Therefore, we tested if wheat *VTL* genes could respond at the transcript level when subjected to changing Fe concentration. The expression in roots and shoots of wheat seedlings was measured after subjecting them for 3 and 6 days of starvation. Our expression analysis suggests that in roots all the *VTL* genes (*TaVTL1*, *TaVTL2*, *TaVTL4* and *TaVTL5*) were downregulated at both the days, whereas, only *TaVTL5* was upregulated at 6 days of starvation (Figure 4A).

**Figure 4:**
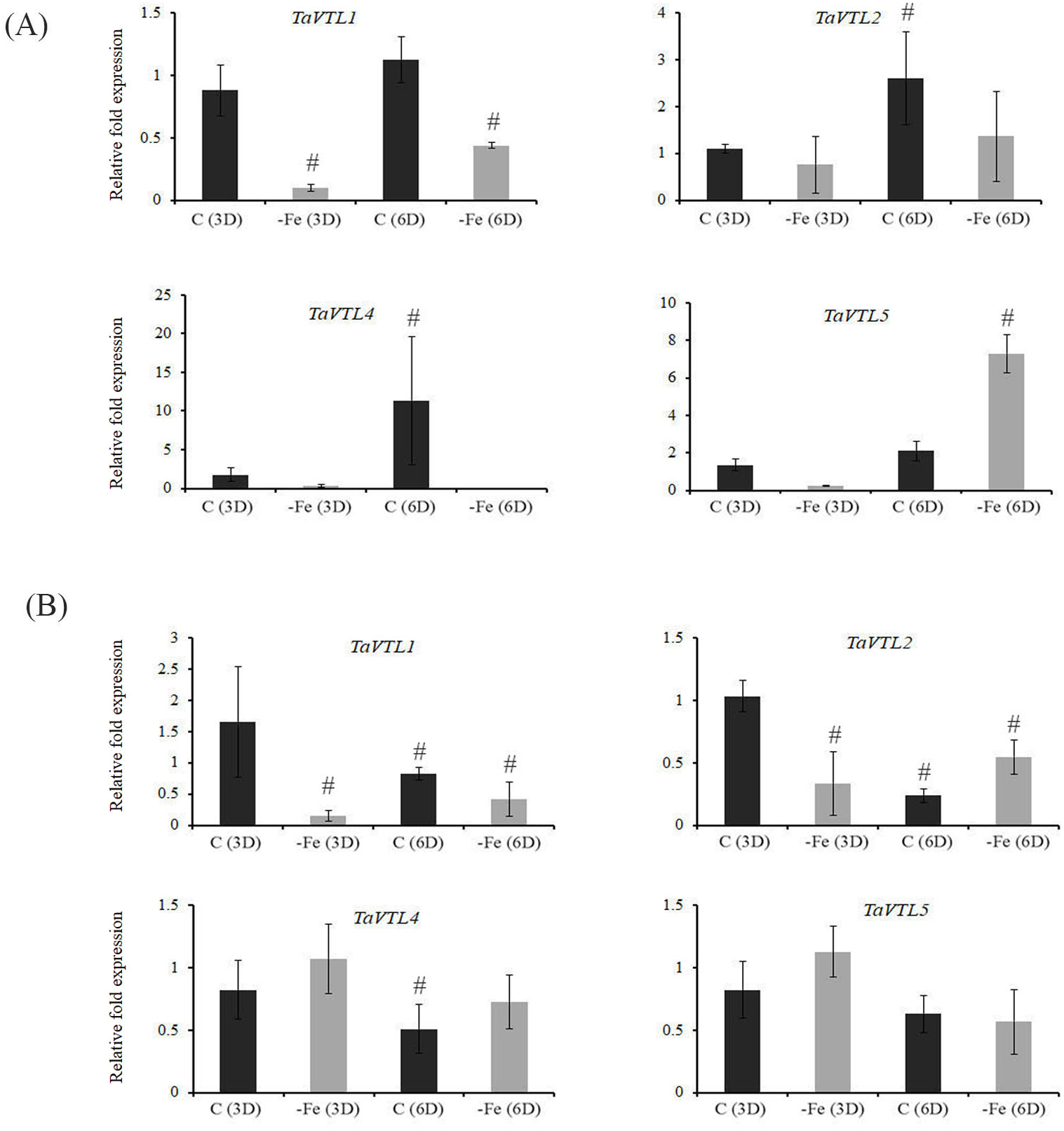
Tissue-specific qRT-PCR expression analysis of wheat VTL genes during Fe deficiency (−Fe) and in the control (C) conditions. Wheat seedlings were subjected to Fe deficiency for 3 and 6 days represented as −Fe(3D) and −Fe(6D). The controls for the respective time points are represented as C(3D) and C(6D). (A) Fold expression analysis was performed in roots and (B) in shoots. 2 μg of total RNA was used for the cDNA preparation. Relative fold expression levels were calculated relative to C(3D). C_t_ values were normalized using wheat *ARF1* as an internal control. Vertical bars represent the standard deviation. # represents the significant difference at p < 0.05 with respect to their respective control treatments.

Similarly, in shoots also all the expression of wheat *VTL* genes was suppressed except for *TaVTL2* that was upregulated only on 6 days post starvation (Figure 4B). These expression data demonstrate that under Fe deprivation *VTL* gene expression are negatively regulated in wheat seedling. Transcriptomic sequencing data from wheat seedlings after 20 days of Fe starvation (SRP189420) was also used to check expression response upon Fe starvation. Categorically, *TaVTL5* group genes were seen to be upregulated upto 12-fold, with *TaVTL5-2B_6* showing upregulation of ~60 fold, although the expression was not very high (Figure S3).

Next, we performed the gene expression analysis under the excess Fe regime. This was done to test if wheat *VTL* genes could be potentially involved in detoxification of excess Fe. Interestingly, we observed a significant up-regulation of all the *TaVTL* genes in roots at both the time points. Out of all, *TaVTL4* showed the highest fold gene expression (~100 fold) when compared to its control (Figure 5A). *TaVTL2* show very early and high expression response, whereas both *TaVTL1* and *TaVTL5* were highly expressed at 6 days of treatment. At this time their gene expression level was more than ~14 fold compared to control. In contrast, in shoots most of the wheat *VTL* genes were expressed at the 3 days of treatment with *TaVTL1* and *TaVTL2* showing the transcript accumulation of 8-14 folds with respect to their control (Figure 5B).

**Figure 5:**
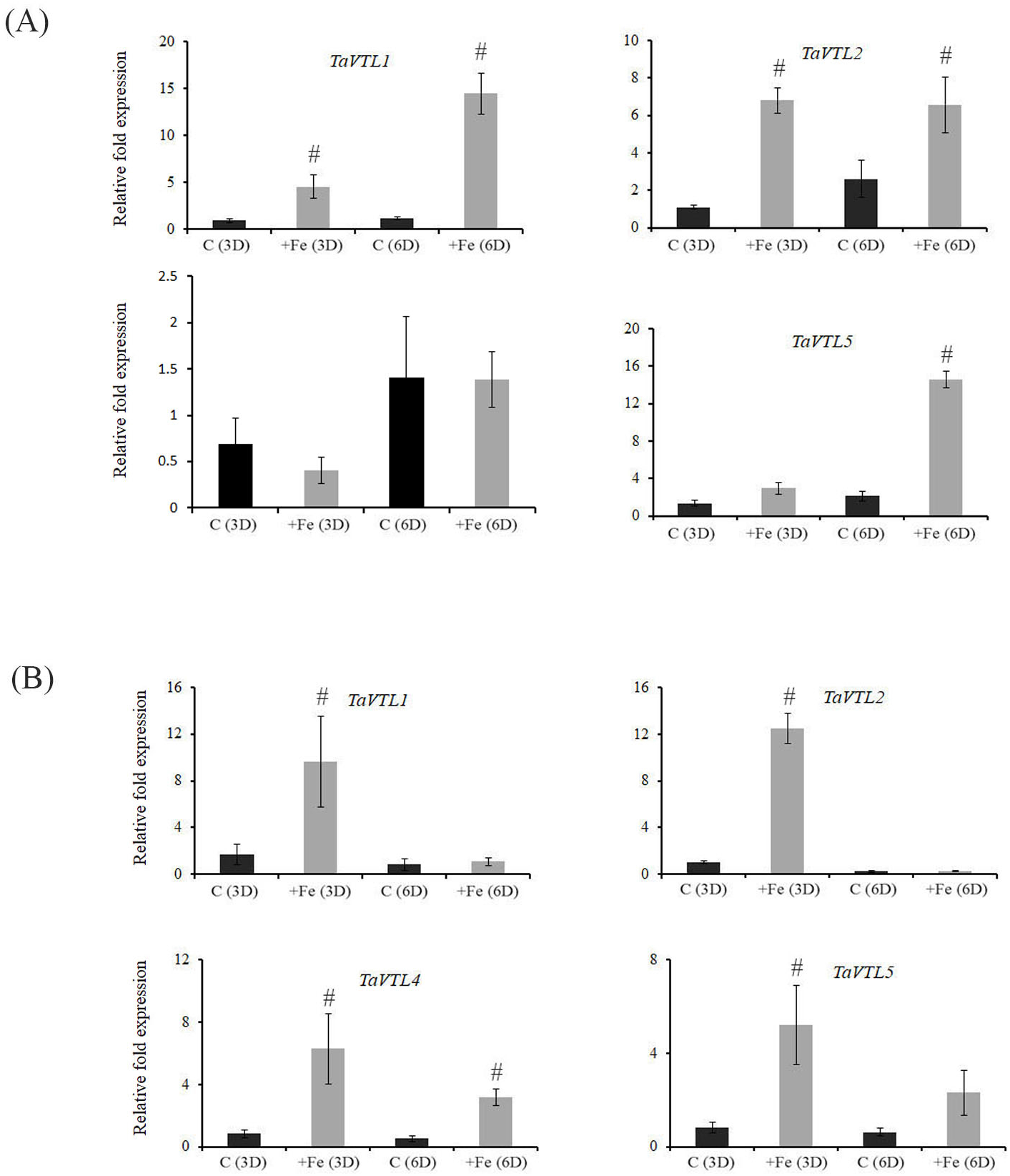
Tissue-specific qRT-PCR expression analysis of wheat VTL genes during Fe surplus (+Fe) and in the control (C) conditions. Wheat seedlings were subjected to Fe surplus for 3 and 6 days represented as +Fe(3D) and +Fe(6D). The controls for the respective time points are represented as C(3D) and C(6D). (A) Fold expression analysis was performed in roots and (B) in shoots. 2 μg of total RNA was used for the cDNA preparation and relative fold expression levels were calculated relative to C(3D). C_t_ values were normalized using wheat *ARF1* as an internal control. Vertical bars represent the standard deviation. # represents the significant difference at p < 0.05 with respect to their respective control treatments.

### 2.5 Manganese, Zinc and Copper deficiency causes differential changes in VTL expression

Wheat *VTL* genes showed high similarity to previously known *VIT* genes. In addition to Fe, *VIT* genes are known to be affected by the perturbed concentration of Zn and Mn [15]. Since many of these cation transporters are known for their reduced substrate specificity, therefore, expression of wheat *VTL* genes during Zn and Mn deprivation was also studied. In general, during the changing regimes of Zn and Mn, wheat *VTL* genes showed specific expression in a tissue-specific manner (Figure 6). *TaVTL2* was the only gene showing enhanced accumulation of its transcript under Zn deficiency in both root and shoot tissue, whereas *TaVTL1* and *TaVTL5* showed high expression in roots only under Zn deficiency. No significant changes in the expression of *TaVTL4* were observed for the studied time point under the changing Zn regime. In contrast, no induction of wheat *VTL* genes was observed in roots under Mn deficiency with respect to its control, whereas, in shoots, *TaVTL2* and *TaVTL4* showed high transcript accumulation. Under Cu deficiency, all *VTL* genes showed an induced expression in shoots while only two of the genes, including *TaVTL1* and *TaVTL2* were upregulated in roots.

**Figure 6:**
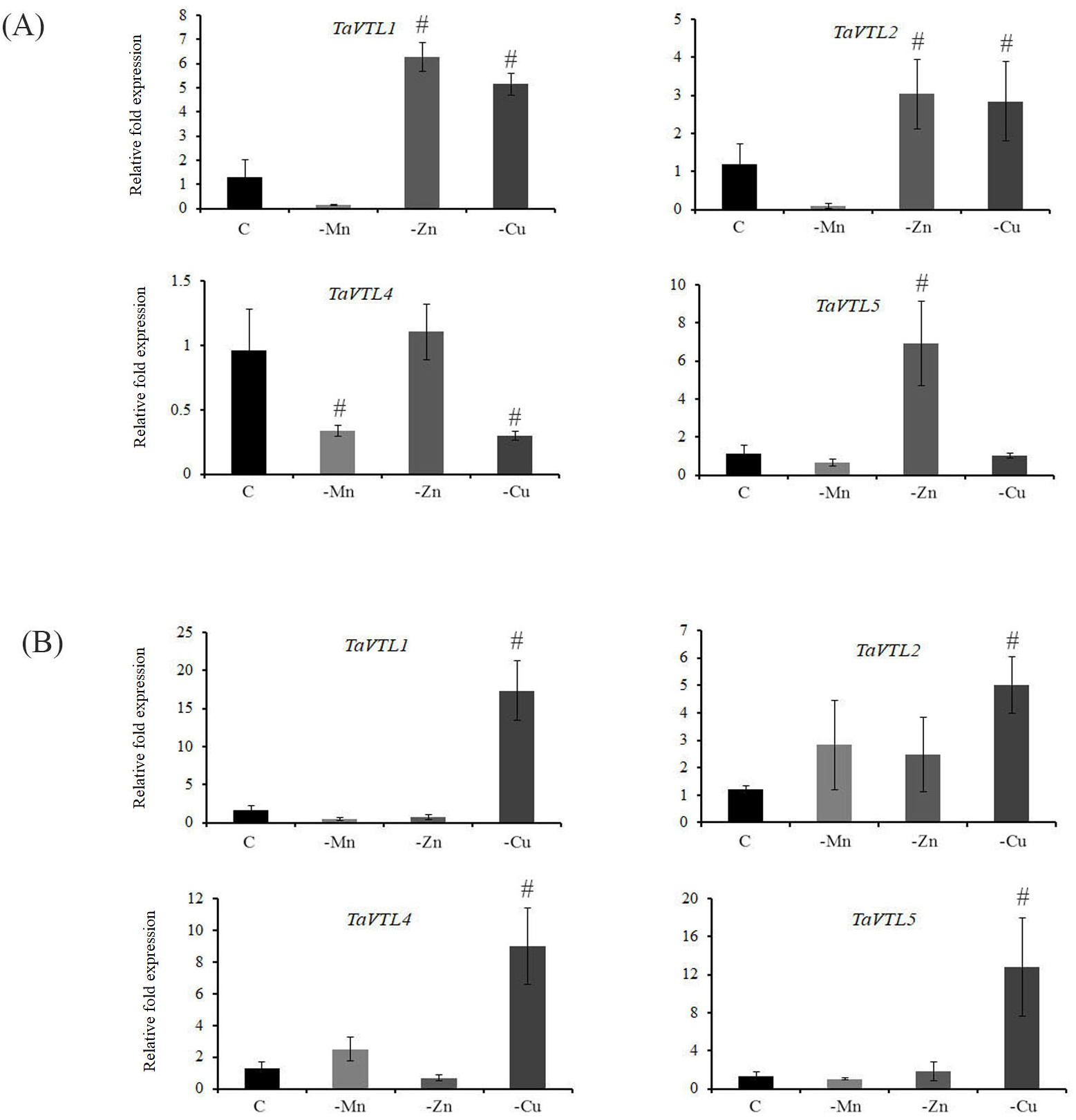
Tissue-specific qRT-PCR expression analysis of wheat VTL genes during Mn (−Mn), Zn (−Zn) and Cu (−Cu) deficiency with rest to the control (C) conditions. (A) Fold expression analysis was performed in roots and (B) in shoots. 2 μg of total RNA was used for the cDNA preparation and relative fold expression levels were calculated relative to control tissue (C). The C_t_ values were normalized using wheat *ARF1* as an internal control. Vertical bars represent the standard deviation. # represents the significant difference at p < 0.05 with respect to Control tissue.

### 2.6 Heavy metal (Ni, Cd and Co) mediated expression of VTL genes

To check the effect of the heavy metal on the gene expression pattern, wheat seedlings were subjected to treatment with Ni, Cd and Co and; expression of *VTL* genes was performed. In the treated plants, decreased growth of the shoot and root length was observed, suggesting that heavy metals could affect the plant performance (data not shown). Interestingly, none of the wheat *VTL* genes showed enhanced expression in roots after 15 days of heavy metal exposure. In shoots, only under the metal Co-treatment *TaVTL2*, *TaVTL4* and *TaVTL5* show significantly higher expression as compared to control shoot samples (Figure 7). This data suggested the metal-specific expression of wheat *VTL* genes in a tissue-specific manner. The previously reported wheat *VIT* genes showed grain specific expression data. Surprisingly, *VTL* genes showed very low or no expression in grains or their tissue parts, suggesting their probable roles in the organs of the plants (Figure S4).

**Figure 7:**
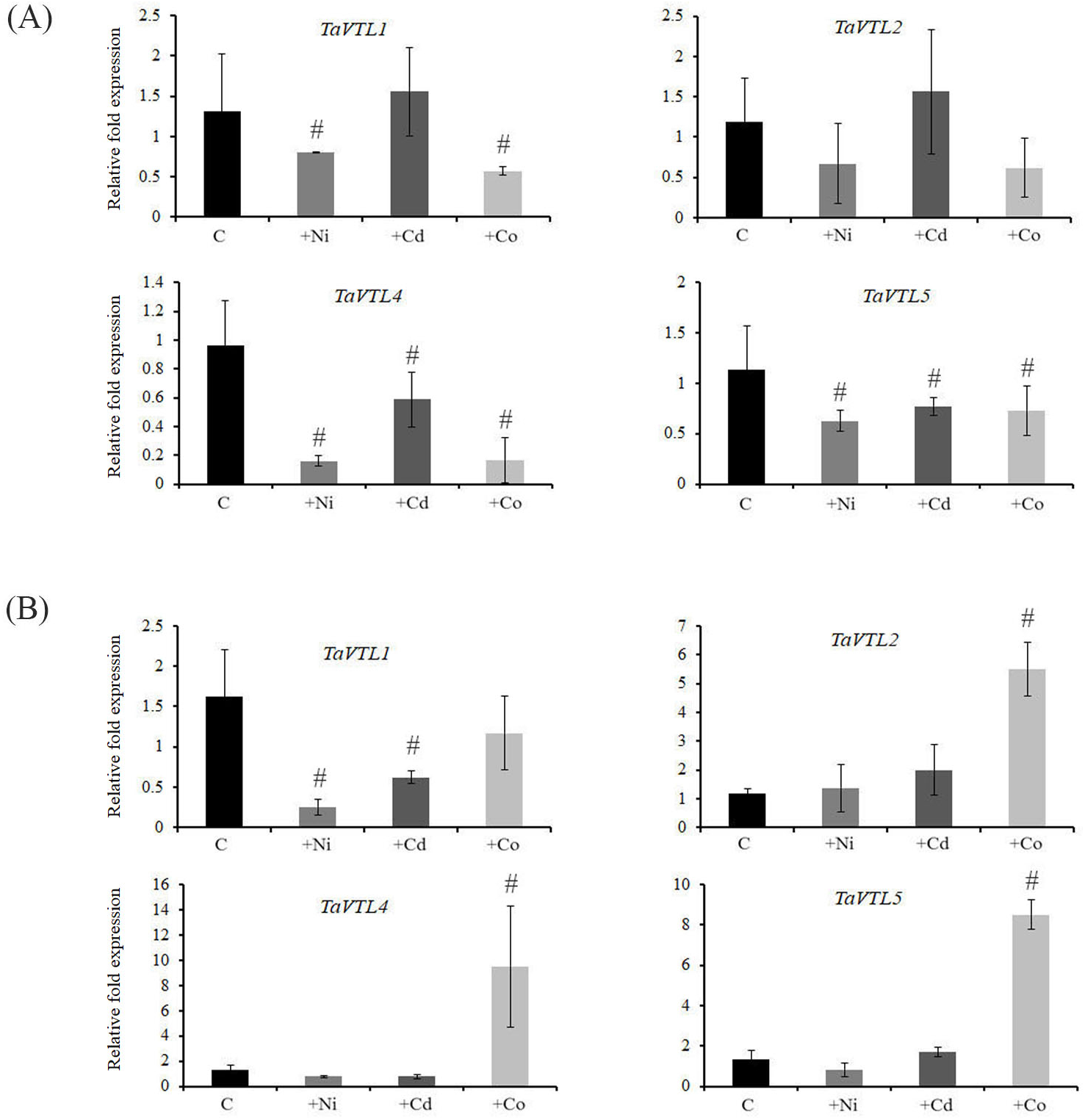
Tissue-specific qRT-PCR expression analysis of wheat VTL genes upon heavy metal treatments. Wheat seedlings were exposed to Ni (+Ni,50 μm), Cd (+Cd, 50 μm) and Co (+Co, 50 μm). Control seedlings (C) without any exposure to heavy metals were compared with the treated ones. (A) Fold expression analysis was performed in roots and (B) in shoots. 2 μg of total RNA was used for the cDNA preparation and relative fold expression levels were calculated relative to control samples. Ct values were normalized using wheat *ARF1* as an internal control. Vertical bars represent the standard deviation. # represents the significant difference at p < 0.05 with respect to their respective control treatments.

## 3. Discussion

Fluctuation in the nutrient condition in the soil results in drastic adaptions by the plants. In general, plants rely on different physiological and molecular processes to minimize nutrient stress [28]. In this regard, MFS gene family plays an important role to provide the tolerance as well as mobilization of important minerals, including micronutrient to the foliar parts including seeds [17]. In this study, the characterization of the new gene family VTL was done. Our data reinforce the importance of this gene family for their roles not limited during metal homeostasis but also a good potential for biofortification in cereal crops such as wheat and rice.

MFS family has been widely explored for its role as metal transporters and providing the necessary support to provide multiple functions in plants [29]. Previously, five *VTL* genes were reported in *Arabidopsis* and five genes were reported in rice for this sub-class. Our study in wheat resulted in the identification of a maximum number of *VTL* genes from any crop plants. The high number of genes is due to the presence of multiple homoeologues and occurrence of the duplication of many wheat *VTL* genes. Interestingly, chromosome 2 has been linked with multiple quantitative trait loci (QTL) for the high grain Fe and Zn [30]. Further dissection is required in this direction to identify if any of the wheat VTL could be linked with the loading of micronutrient in grains. Based on our expression analysis and the support from the previous studies, it could be suggested that *VTL* genes could also be involved in providing the tolerance to high levels of Fe and Zn in the soils [25]. In fact, the predicted localization data indicate that VTL could be localized at either the plasma membrane or the vacuolar membrane. AtVTL1 was reported to be localized in the vacuolar membrane and others also be associated with the plasma membrane [25]. Our PSORT analysis suggests that most of the wheat VTL are localized in the vacuolar membrane, thus making them a suitable candidate for sequestering micronutrients such as Fe and Zn.

The substrate specificity of the metal transporters is a major bottleneck to achieve high Fe and Zn rich grains. Manipulating the specificity of these metal transporters to enrich the Fe and Zn remains the major challenge [20]. Therefore, studying the expression pattern of *VTL* genes in the presence of heavy metals could provide preliminary clues to employ such strategies. Consequently, the study was undertaken to see the influence of other metals like Ni, Cd and Co. The expression of wheat *VTL* genes in roots and shoots suggested an interesting phenomenon, where no significant changes in the expression of their transcript was observed when exposed to either Ni and Cd. In contrast, only Co was able to induce the expression of *TaVTL1*, *TaVTL2*, *TaVTL4* and *TaVTL5* in shoots only. This data suggests the controlled expression of wheat *VTL* genes in specific tissues. Additionally, besides Fe homeostasis, the vacuolar transporters are also linked with the impaired activity of Zn and Mn transport [31]. In our study only, *TaVTL2* was significantly induced by the Mn deficiency in shoots. No such effects were observed in roots wherein, all the quantified wheat VTL showed downregulation under Mn deficiency. Interestingly, *TaVTL1* and *TaVTL2* showed upregulation in roots under Zn deficiency. The tissue dependent expression patterns of wheat *VTL* genes under the changing regimes of the metal exposure suggest was observed. It has been observed that *VTL* genes from *Arabidopsis* showed transcriptional changes in response to Fe, Zn and Mn [25]. Based on the previous work and our results it could be suggested that regulation of the VTL genes at the transcript level could be conserved. Our works gene expression data also corelates with the presence of multiple cis-elements in the promoter of wheat VTL genes.

Herein, a detailed inventory, structure and expression characterization of wheat *VTL* genes was performed. The work presented here provides the preliminary clue for the expression characterization of wheat *VTL* genes that further remains to be characterized for their *in-planta* functional activity using forward and reverse genetics approaches.

## 4. Conclusions

The present work lead to the identification of high number of VTL genes from hexaploid wheat. Because of polyploidization, a very high number of genes from this sub-family was identified. The presence of high number of VTL been restricted to only chromosome 2, 4 and 6 of the wheat genomes. The expression of these gene under metal stress including changes in the presence of Fe and Zn concentrations and exposure to heavy metals reinforce the importance of this gene-family during metal homeostasis. Therefore, VTL class of gene family in wheat requires further characterization for their localization patterns and for their functional activity.

## 5. Materials and Methods

### 5.1 Identification of VIT family and classification of VTL genes in wheat

For the identification of wheat *VTL* genes, the Ensembl database was used to extract *VIT* family genes (Pfam ID: PF01988) for wheat. The identification was confirmed by bidirectional blast analysis. VIT family sequences from Arabidopsis, rice, maize, *Brachypodium* were also extracted using Pfam search. The identity of VIT family genes was further validated by confirming the presence of CCC1-like superfamily domain using NCBI-CDD domain search. CCC1 sequence for *S. cerevisiae* was also retrieved from its genome database. To separate out *VTL* genes from *VIT* genes and for further phylogenetic analysis, all the proteins were aligned through MUSCLE alignment and an unrooted neighbor-joining phylogenetic tree with 1000 bootstrap replicates was constructed with all the retrieved sequences. The tree was constructed through MEGA-7 [32]. Rice vacuolar iron transporter homolog 1-5 from UniProt were used for the nomenclature of the twenty-three *TaVTL* sequences based on the closest orthologs. The naming of the genes indicates the chromosome number and the sub-genome on which they are present.

### 5.2 Conserved domains and motif detection, analysis of gene, promoter and protein structure

Wheat VIT family genes were searched for conserved domains using NCBI-CDD database[33]. MEME suite v5.1.0 was used for further analysis to identify the common conserved motifs for both VIT and VTL proteins. The maximum number of motifs was set to 15 for MEME analysis. Gene structure for VITs and VTLs were studied using (GSDS) (http://gsds.cbi.pku.edu.cn/) [34] using genomic and CDS sequences. Sub-cellular localization and TM domains were predicted using web-based prediction programs Wolf PSORT and Phobius respectively[35]. For promoter analysis, ~2Kb promoter elements of the corresponding wheat VTL genes were surveyed for the presence of the respective cis-elements. The promoter sequence was obtained for the respective genes suing the IWGSC

### 5.3 Plant materials and growth conditions

For stress experiments, hexaploid wheat *Triticum aestivum* cv. C-306 (received from Punjab Agriculture University, Ludhiana) was used. Briefly, seeds were surface sterilized using 1.2% sodium hypochlorite prepared in 10% ethanol and then rinsed twice with autoclaved MQ. The seeds were kept on moist filter paper inside a Petri dish and stratified for 1 day at 4 °C in dark condition. Stratified seeds were further kept for germination for 6 days at room temperature. The remaining seed/endosperm was excised from seedlings at one leaf stage and was shifted to phytaboxes (10-12 seedling / phytabox) containing the Hoagland nutrient media for respective treatments. The standard composition of nutrient media for control includes 6 mM KNO3, 1 mM MgSO_4_, 2 mM Ca(NO3), 2mM NH_4_H_2_2PO_4_, 20 μM Fe-EDTA, 25μM H3BO3, 2 μM MnSO4, 0.5 μM CuSO4, 2 μM ZnSO4, 50 μM KCl and 0.5 μM Na2MoO4. The variable concentrations used for treatments were excess Fe (+Fe; 200μm), Fe starvation (−Fe;2μm), Zn deficiency (−ZnSO4; 0μm), Mn deficiency (−MnSO4; 0μm), Cu deficiency (−CuSO4; 0μm), Cadmium stress (+Cd; 50 μm), Cobalt stress (+Co; 50 μm) and Nickel stress (+Ni; 50 μm). The aerobic condition was provided in hydroponics and the media was replaced every alternate day to avoid any contamination and drastic nutrient depletion. The respective roots and shoots samples belonging to iron deficient and sufficient plant groups were collected at 3 and 6 days after stress (D). For the rest of the treatments, root and shoot, samples were collected on the 15^th^ Day of treatments. All the experiments were performed in a growth chamber under controlled environmental conditions at 22-24 °C temperature, 65-70% humidity, at a photoperiod of 16 hours day and 8 hours night and 300 nm of light.

### 5.4 RNA isolation and cDNA preparation

The collected root and shoot samples were ground separately in liquid nitrogen. Total RNA from respective samples was extracted by TRIZOL based method. The extraction was followed by the DNase treatment using Turbo DNAfree kit (Invitrogen, USA) to remove any genomic DNA contamination in the RNA samples. Subsequently, RNA purity was checked and quantified for the preparation of the cDNA. 2 μg of total RNA was used for cDNA synthesis using superScript III First-Strand Synthesis System (Invitrogen, USA). The cDNA quality was ascertained by using internal control and was further diluted 20X and used for gene expression studies.

### 5.6 Quantitative-real time PCR (qRT-PCR) expression analysis

To perform quantitative real time-PCR (qRT-PCR), forward and reverse primers of *TaVTL* genes were designed and used as listed in Table S5. The primers were designed from the conserved region of the all homoeologs of each gene. For *TaVTL5* the primers were designed from the conserved region of 9 sequences, the significant conserved region was not found for remaining 4 homoeologs (TraesCS2B02G454900, TraesCS2B02G610400, TraesCS2D02G431900, TraesCS2D02G588000). qRT-PCR was performed in 7500 Real-Time PCR System (Applied Biosystems, USA) using 1/20 times dilution of the respective cDNAs. All qRT-PCR reactions were performed using SYBR Green I (QuantiFast® SYBR® Green PCR Kit, Qiagen, Germany) chemistry and ARF (ADP-Ribosylation Factor: *TaARF1*—AB050957.1) as an internal control [36]. The efficiency of the qRT-PCR was checked and melt curve analysis was performed for each of the PCR reactions as per the guidelines. Gene expression analyses was carried out with three biological replicates and two-three technical replicates. Relative fold expression of genes was determined based on delta-delta CT-method (2^−ΔΔCT^) [37].

### 5.7 RNA-seq expression analysis for VIT family genes

To get the transcript expression levels for VIT family genes under Fe stress, RNAseq data from SRA project ID SRP189420 was utilized to extract transcript expression values (as FPKM) from control as well as Fe starved wheat root samples using the cufflinks pipeline. Subsequently, for expression analysis of *VTL* and *VIT* genes in wheat grain tissue developmental time course [38], expression values as Transcripts Per Kilobase Million (TPM) were retrieved from expVIP database. Expression values from both studies were then used to plot heatmaps using MeV software.

### 5.8 Statistical Analysis

Excel was used for data analysis. The mean values were calculated form the standard deviation including three technical replicates form at least three biological replicates. Student t-tests were used to observe the significant differences between the mean values of treatment and control plants. The significance threshold used was ^#^*P* < 0.05.

## Acknowledgments

All the authors thank Executive Director, NABI for facilities and support. This research was funded by the NABI-CORE grant to AKP. Support from International Wheat Genome Sequencing Consortium for providing the high-quality wheat genome resources is highly appreciated.

## Author Contributions

Conceptualization, SS., AKP; methodology, SS, AK, VM and AKP; formal analysis, SS, AKP, GK and HR; investigation, SS, VM; writing—original draft preparation, AKP., AK., JK; writing—review and editing, AKP., AK., GK., and HR; visualization, AKP; funding acquisition, AKP.

