## Supplementary Figures S1 to S4 for "Gene expression pattern of vacuolar-iron transporter-like (VTL) genes in hexaploid wheat during metal stress"

### Slide 1
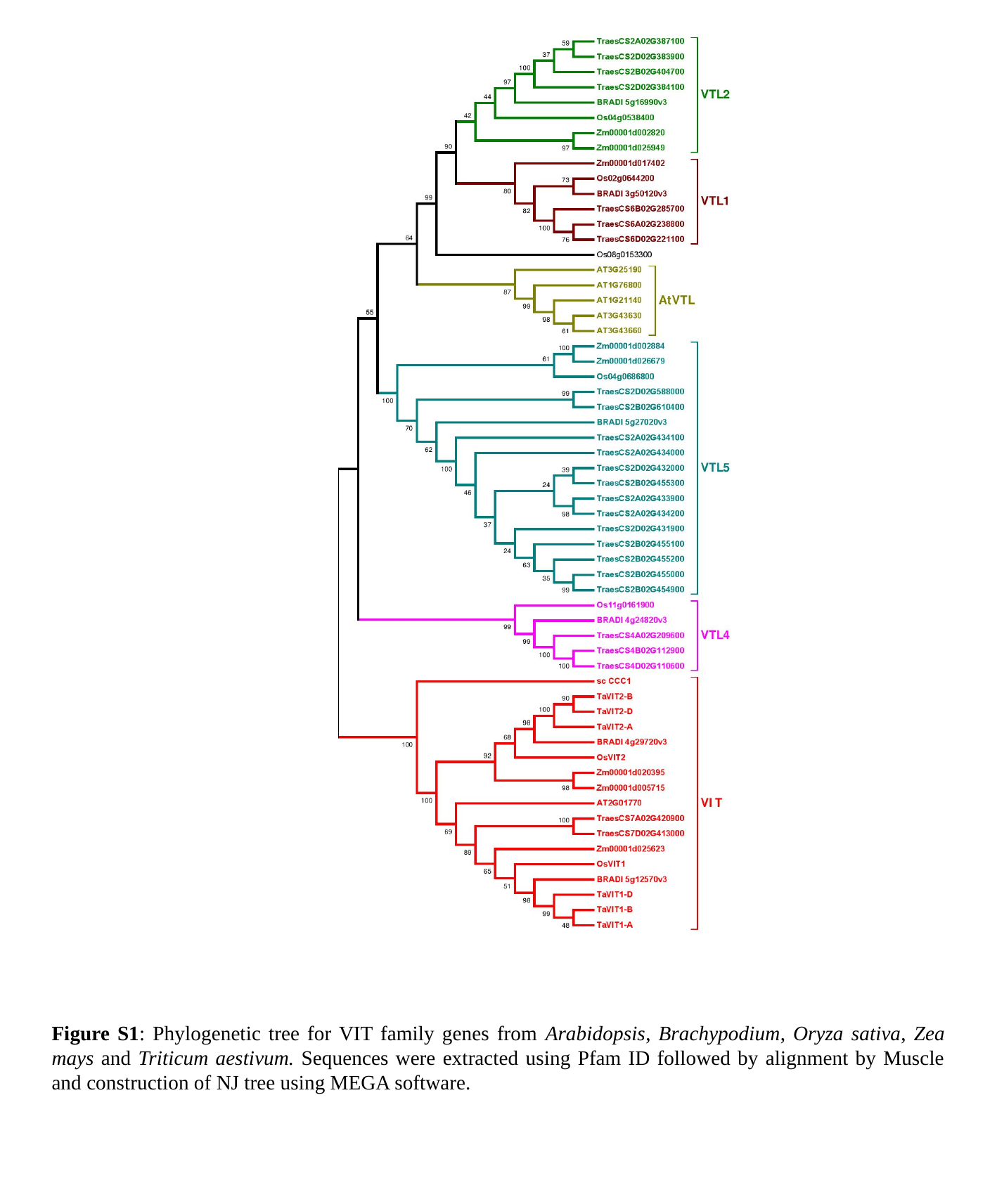

Figure S1: Phylogenetic tree for VIT family genes from Arabidopsis, Brachypodium, Oryza sativa, Zea mays and Triticum aestivum. Sequences were extracted using Pfam ID followed by alignment by Muscle and construction of NJ tree using MEGA software.

### Slide 2
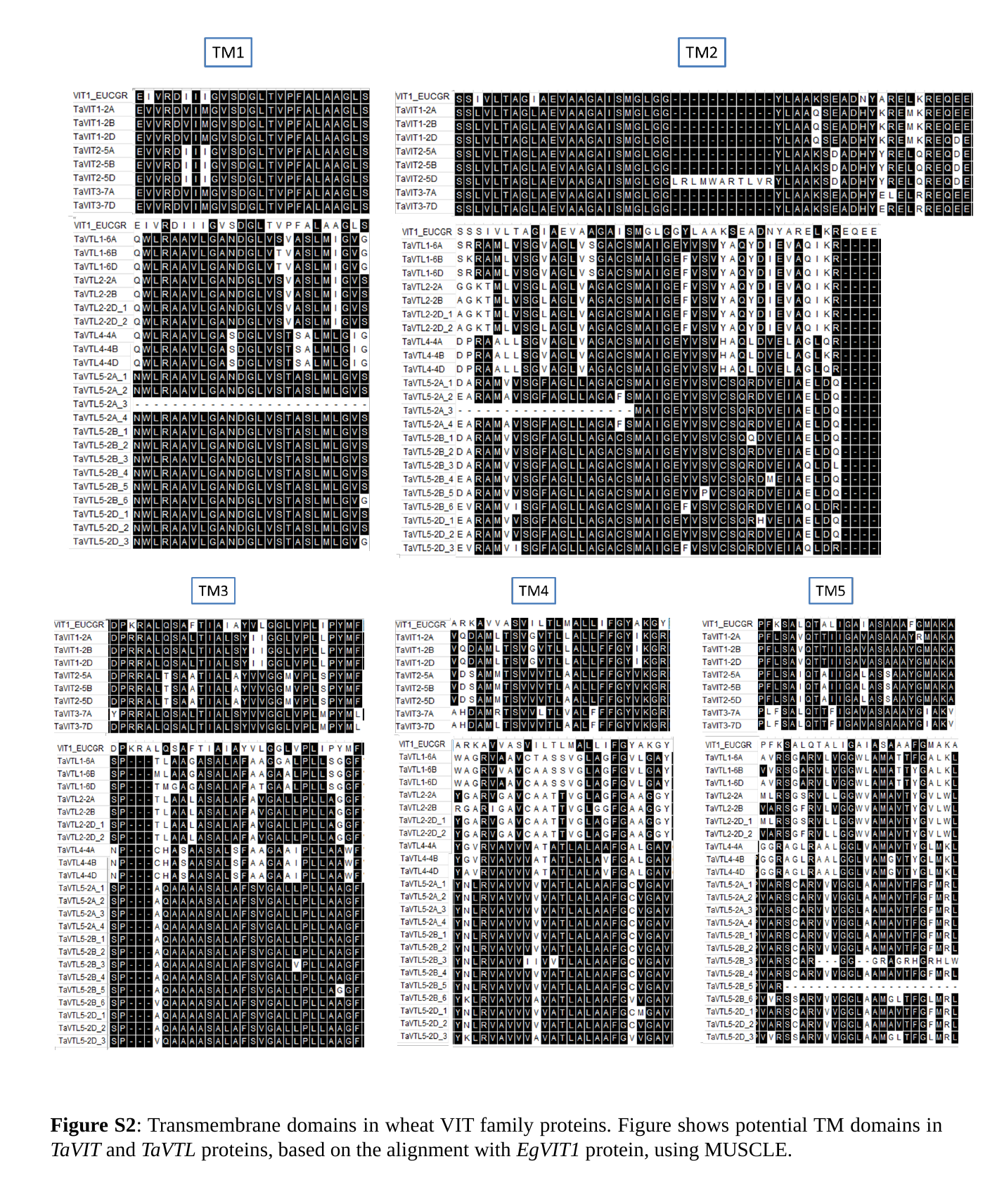

Figure S2: Transmembrane domains in wheat VIT family proteins. Figure shows potential TM domains in TaVIT and TaVTL proteins, based on the alignment with EgVIT1 protein, using MUSCLE.

### Slide 3
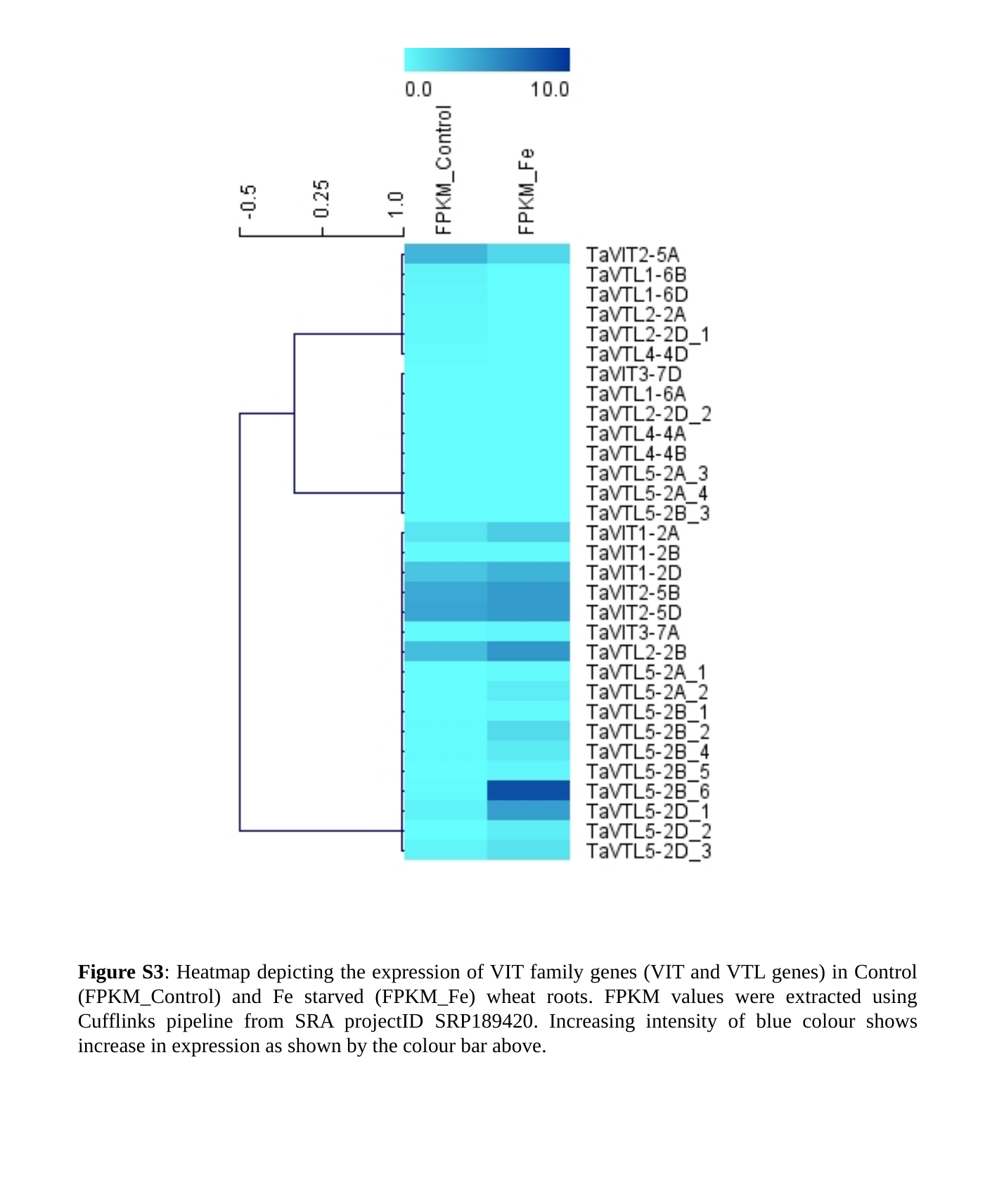

Figure S3: Heatmap depicting the expression of VIT family genes (VIT and VTL genes) in Control (FPKM_Control) and Fe starved (FPKM_Fe) wheat roots. FPKM values were extracted using Cufflinks pipeline from SRA projectID SRP189420. Increasing intensity of blue colour shows increase in expression as shown by the colour bar above.

### Slide 4
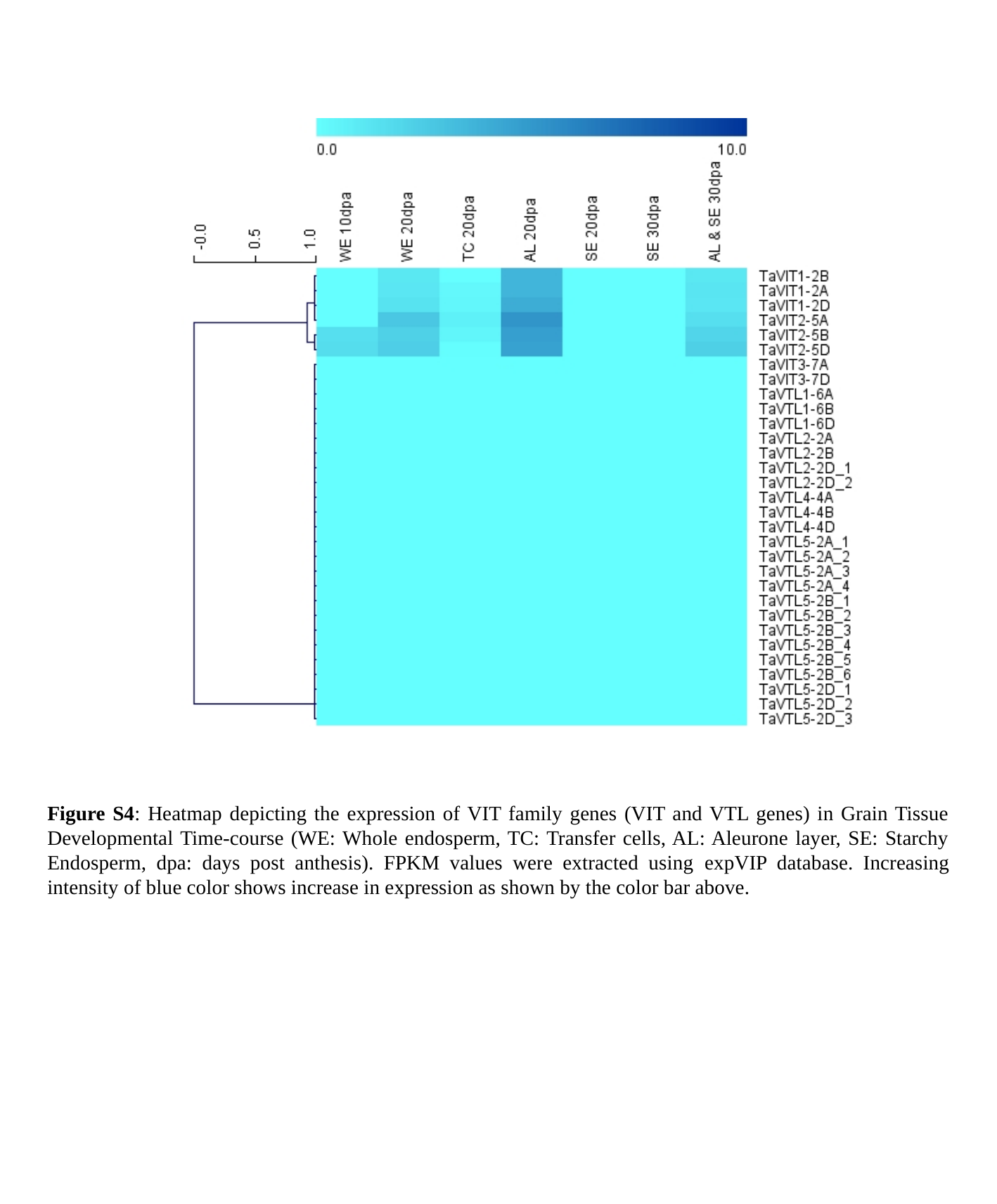

Figure S4: Heatmap depicting the expression of VIT family genes (VIT and VTL genes) in Grain Tissue Developmental Time-course (WE: Whole endosperm, TC: Transfer cells, AL: Aleurone layer, SE: Starchy Endosperm, dpa: days post anthesis). FPKM values were extracted using expVIP database. Increasing intensity of blue color shows increase in expression as shown by the color bar above.
