## Supplementary Table S5 for "Gene expression pattern of vacuolar-iron transporter-like (VTL) genes in hexaploid wheat during metal stress"

**Supplementary Table S5:** List of gene specific primers used for qRT-PCR for *TaVTL genes.*

| **Gene name** | **Amplicon size** | **Primer sequence (5'-3')** |
| --- | --- | --- |
| TaVTL5 F | 151bp | AGCTGGACCAGGCCGGAAAG |
| TaVTL5 R | 151bp | CACGACGACCACCACGGC |
| TaVTL1 F | 120bp | TCATGATCGGCGTCGGCGCC |
| TaVTL1 R | 120bp | CGGCGGGTGCGCTTGATCTG |
| TaVTL4 F | 179bp | ACCGACAATGACACCAAGCTCGCT |
| TaVTL4 R | 179bp | GACGGCGGCGCGCAGCCAC |
| TaVTL2 F | 203bp | ACATGGCCCGCGCGCAGTGG |
| TaVTL 2 R | 203bp | GATGTCGTACTGCGCGTACACGG |
